# Dissecting the Ca^2+^ dependence of *Mycobacterium tuberculosis* DesA1 function

**DOI:** 10.1101/2023.03.31.535087

**Authors:** Mamata Savanagouder, Ravi Prasad Mukku, Uday Kiran, Chaitanya Veena Yeruva, Nandhini Nagarajan, Yogendra Sharma, Tirumalai R. Raghunand

**Author notes:** Correspondence and reprints Tirumalai R. Raghunand -. Equal contribution (Mamata Savanagouder, Ravi Prasad Mukku). Email information: Mamata Savanagouder - Ravi Prasad Mukku - Uday Kiran - Chaitanya Veena Yeruva - Nandhini Nagarajan - Yogendra Sharma. Institute of Animal Reproduction and Food Research, Polish Academy of Sciences in Olsztyn, Tuwima 10 Str,10-748 Olsztyn, Poland (Mamata Savanagouder). Sanofi Healthcare India Pvt. Ltd., Athvelly, Medchal, Hyderabad, 501401, India. The Clinic’les Pvt. Ltd., Meyyanur, Salem 636004, India. IISER Berhampur (Main Campus) Vigyanpuri, At/Po: -Laudigam, Konisi, Dist. Ganjam, Odisha 760003, India.

## Abstract

*Mycobacterium tuberculosis (M. tb)* has a complex cell wall, largely composed of mycolic acids and long-chain fatty acids that play a crucial role in maintaining its integrity and permeability. This complex lipid structure has a role in abrogating the process of phagosome-lysosome fusion and infection establishment. The *M. tb* desaturase A1 (DesA1) catalyzes the introduction of position-specific double bonds, a key step in the biosynthesis of a diverse range of mycolic acids. We have previously demonstrated that *M. tb* DesA1 is a Ca^2+^-binding protein, belonging to the extended βγ-crystallin superfamily. Using a combination of biophysical and genetic approaches, we investigated the structural and functional significance of Ca^2+^ binding on DesA1 activity. A protein unfolding assay of the protein in the presence and absence of Ca^2+^ shows that Ca^2+^ binding imparts structural stability to DesA1. To identify the role of Ca^2+^, we introduced mutations at key residues in the identified Ca^2+^-binding motif of DesA1 and generated F303A, E304Q, and F303A-E304Q variants of DesA1. We identified F303 as a hot point which disables the protein for Ca^2+^ binding. Two other mutations E304Q and F303A-E304Q showed reduced Ca^2+^ binding. Complementation of a conditionally complemented *desA1* deletion mutant strain of *Mycobacterium smegmatis* with these mutants, either failed to complement its growth phenotype or led to a compromise in complementation. In addition, the F303A and F303A-E304Q complements exhibit increased sensitivity to isoniazid, a first-line anti-tubercular drug, pointing to a cell wall permeability defect in these strains. Our findings highlight the critical importance of Ca^2+^ in the functioning of DesA1 and its implicit role in the maintenance of mycobacterial cellular integrity.

## INTRODUCTION

The lipid-rich cell wall of *Mycobacterium tuberculosis* (*M. tb*) - the causative agent of tuberculosis (TB), contains a unique class of fatty acids called mycolic acids (MA). These are α-alkyl, β-hydroxyl long-chain fatty acids consisting of a long mero chain (C_40_ - C_60_) and a short a-chain (C_22_ – C_26_), further addition of various functional groups such as double bonds, keto, ester, epoxy, methoxy, and cyclopropane rings gives rise to diverse MAs. MA biosynthesis involves two fatty acid synthase (FAS) complexes: FAS-I and FAS-II, which synthesize the short a-chain and the long mero chain, respectively. This is followed by the introduction of double bonds by desaturases in the mero chain, an essential step for the addition of various functional groups to generate diverse MAs. The modified mero chain further undergoes Claisen condensation with the α-chain to yield the characteristic α-alkyl, β-hydroxyl MAs. MAs are mainly found covalently linked to arabinogalactan and thereby to the peptidoglycan to form the mycolic acid-arabinogalactan-peptidoglycan complex (MAPc). They are also found in the outer membrane as free mycolates such as glucose monomycolate, trehalose monomycolate, and trehalose dimycolate - also known as the ‘cord factor’. MAs contribute to the integrity and low permeability of the mycobacterial cell wall, thereby conferring mycobacteria with intrinsic resistance to commonly used antibiotics. Isoniazid and ethionamide, first- and second-line anti-TB drugs, which target InhA in the MA biosynthesis pathway, have been widely successful in the treatment of TB [1]. With the rise of multidrug and extensively drug-resistant *M. tb* strains, there is an urgent need for identifying potential drug targets and alternate treatment strategies. Given the importance of MA in cell viability and the pathogenesis of *M. tb*, the MA biosynthesis pathway is an attractive target for the development of novel anti-TB drugs.

The *M. tb* genome encodes three putative aerobic desaturases encoded by *desA1, desA2, and desA3* [2]. Of these, DesA3 (Rv3229c) is involved in oleic acid biosynthesis [3] while DesA1 (Rv0824c, https://mycobrowser.epfl.ch/genes/Rv0824c) and DesA2 (Rv1094, https://mycobrowser.epfl.ch/genes/Rv1094) are annotated as acyl-ACP desaturases. Structural analysis of DesA2 revealed that it is a homodimer and is related to plant fatty acid desaturases [4], and a recent report uncovered an early role for DesA2 in the mycolic acid biosynthesis machinery [5]. A homologue of *M. tb desA1, MSMEG5773* (*desA1)* is an essential gene in *M. smegmatis*, and a conditional mutant showed loss of MA synthesis and cell viability under non-permissive conditions. Interestingly, episomal expression of *M. tb desA1* in this mutant rescued cell viability [6]. Studies have shown that *desA1/2* are upregulated during hypoxia-induced in culture [7] and in macrophages, leading to the hypothesis that during early events of infection, increased desaturation events allow *M. tb* to tweak mero-chain modifications towards a phenotype that favors the establishment of infection [8]. The essentiality of *desA1* and its role in the pathogenic process therefore makes it an ideal target for drug development.

We have previously identified that DesA1 is a low to moderate affinity Ca^2+^-binding protein that contains βγ-crystallin type Greek key signatures [9]. As Ca^2+^ binding is shown to impart structural stability to microbial homologues of βγ-crystallins [10, 11], we identified hot points by mutating two residues - Phe 303 and Glu 304, proposed to participate in Ca^2+^ binding to DesA1, and assessed their role on the structure and function of DesA1. Also, based on previously observed changes in surface morphology upon expression of *M. tb desA1* in *M. smegmatis* (most likely due to increased mycolic acid synthesis) [9], we hypothesized that DesA1 mutants with reduced or loss of Ca^2+^ binding might cause reduced or defective MA synthesis, leading to a compromised cell wall. To test this possibility, we performed isoniazid sensitivity assays with the mutants as a marker for cell permeability. Our results establish an essential role for Ca^2+^ binding in the structure and function of DesA1, a finding that assumes significance given the indispensable nature of this protein for the viability of *M. tb*.

## MATERIALS AND METHODS

### Bacterial strains, media and growth conditions

*Mycobacterium smegmatis* mc^2^155 and *Escherichia coli* BL21 (DE3) pLysS strains were cultured as described [12]. The following antibiotics or chemicals were added when required - 0.2% acetamide, apramycin (50 µg/ml for *M. smegmatis*), hygromycin (150 µg/ml for *M. smegmatis*), and ampicillin (200 µg/ml for *E. coli*).

### DNA manipulations

Restriction enzymes, T4 DNA ligase, and Q_5_ DNA polymerase were purchased from New England Biolabs. Procedures for DNA manipulations, including polymerase chain reaction, restriction endonuclease digestion, agarose gel electrophoresis, ligation of DNA fragments, cloning, and *E. coli* transformations were carried out as described earlier [13]. DNA fragments used in cloning reactions were purified using the NucleoSpin gel and PCR cleanup kit (Macherey-Nagel) according to the manufacturer’s instructions. Plasmid DNA isolations were performed using the NucleoSpin Plasmid kit (Macherey-Nagel) according to the manufacturer’s instructions.

### Generation of *desA1* mutants

Point mutants *of M. tb desA1 (Rv0824c)* – F303A, E304Q, and F303AE304Q were generated by site-directed mutagenesis using the expression vector pET22b:*desA1*-WT as a template [9]. A two-step PCR amplification was employed to generate *desA1* mutant amplicons using overlapping primers (Table S1) followed by overnight *DpnI* digestion at 37°C. The digested products were transformed into *E. coli* DH5α and selected transformants were processed for plasmid isolation. Mutants were confirmed by sequencing using vector specific primers. For biophysical studies, the mutants were cloned using primers 1-6 (Table S1) between the *Nde1* and *Not1* sites of pET22b generating a C-terminal 6xHis-tagged fusion and transformed into *E. coli* BL21(DE3) pLysS. For complementation and isoniazid sensitivity experiments, the mutants were sub-cloned from pET22b:*desA1*::F303A, pET22b:*desA1*::E304Q, and pET22b:*desA1*::F303A-E304Q using primers 7 & 8 (Table S1) between the *EcoR1* and *BamH1* sites of pMV206A.

### Purification of recombinant proteins

The DesA1 mutants were expressed, purified, and refolded as described for wild type DesA1 [9]. Protein concentrations were estimated using the urea method by absorption at 280 nm in a UV-visible spectrophotometer. For Ca^2+^ binding studies, the purified protein was treated with 10 μM EDTA for 15 min at RT, followed by exhaustive buffer exchange with Chelex-purified buffer (50 mM Tris-HCl pH 8.5, 10 mM KCl).

### Fluorescence and circular dichroism spectroscopy

Intrinsic fluorescence emission spectra of DesA1 and its mutants were recorded on an F-4500 fluorescence spectrofluorimeter (Hitachi Inc., Japan) using 0.1 mg/ml protein in 50 mM Tris-HCl, pH 8.5 and 10 mM KCl. The spectra were recorded from 300 to 450 nm at an excitation wavelength of 295 nm by setting the excitation and emission band passes of 5 nm each. Fluorescence spectra were plotted after subtraction with the appropriate buffer blank using the program Origin 8.0. CD spectra were recorded on a Jasco J-815 spectropolarimeter at various concentrations of Ca^2+^ (10 mM - 1 mM) in 50 mM Tris buffer, pH 7.5, 10 mM KCl. Far-UV CD spectra of DesA1 and its mutants (0.1 mg/ml) were scanned from 190 to 250 nm using a 0.01 cm path length quartz cuvette. Near-UV CD spectra were scanned from 250 to 350 nm at 1 mg/ml protein concentration using a 1 cm path length quartz cuvette. CD spectra were plotted after subtraction with the appropriate buffer blank using the program Origin 8.0.

### Chemical unfolding

Chemical unfolding was performed at increasing concentrations of urea. A stock solution of ultrapure urea (8 M) was prepared in Chelex-purified water and filtered. The concentration of urea was checked on a digital refractometer. Samples were prepared by the addition of either 100 µM EGTA or 1 mM Ca^2+^ to increasing concentrations of urea from 0-7.9 M. Trp emission spectra were recorded on an F-7000 Fluorescence Spectrophotometer (Hitachi Inc, Japan) by exciting the samples at 295 nm. The spectra were recorded from 300-450 nm at 5 nm excitation and emission slits. The unfolding transition data were fit using Origin 8.0, as described earlier [14].

### Isothermal titration calorimetry (ITC)

The energetics of Ca^2+^ binding to the mutants of DesA1 were carried out by ITC at 30°C on a Microcal VP-ITC instrument (Microcal Inc., USA). Protein samples (50 µM in the cell) and 5 mM Ca^2+^ (in the syringe) were prepared in Chelex-purified 50 mM Tris, pH 8.5, 10 mM KCl. The respective blanks were obtained by titrating the buffer with 5 mM Ca^2+^ under identical parameter settings. The data were fit into the available equations using the software Origin 8.0, after subtraction with the appropriate buffer blank.

### Complementation of Δ*MsdesA1* with *M*.*tb desA1* and its mutants

For the complementation assay, WT *desA1* [amplified from genomic DNA of *M. tb* (H37Ra) with gene-specific primers] and its mutants corresponding to F303A, E306Q, and F303AE306Q were cloned into pMV206 (Apra), an *E. coli*-Mycobacterium shuttle vector. A conditional knockout strain, *M. smegmatis* ΔMSMEG5773 (Δ*MsdesA1* - a kind gift from Apoorva Bhatt, University of Birmingham), was used for the complementation assay [6]. The recombinant plasmids were introduced by electroporation into Δ*MsdesA1* and selected on 0.2% acetamide-supplemented 7H10 agar plates containing apramycin and hygromycin. The ability of *M*.*tb* DesA1 and its mutants to complement ΔMSMEG5773 was assessed as previously described [15]. Transformants were cultured in Middlebrook 7H9 (7H9) media with 0.2% acetamide and the requisite antibiotics until they reached log phase, followed by washing of culture pellets with 7H9 media three times to remove the acetamide in the media. To deplete acetamide from the bacterial cytosol, the pellets were resuspended in acetamide free 7H9 media and grown for 12 h. Thereafter, cultures were harvested, dilutions spotted on Middlebrook 7H10 (7H10) agar plates with and without acetamide, and the plates were incubated for 72 h at 37 °C.

### Isoniazid sensitivity assay

To test their sensitivity to isoniazid, recombinant strains of MSMEG5773 expressing WT DesA1, and its F303A, E306Q, and F303AE306Q mutants were grown to log phase, pelleted, washed three times with 7H9 media, and then grown for 12 h in 7H9 media without acetamide. The cells were then harvested, dilutions spotted on 7H10 agar plates containing 4 µg/ml, 8 µg /ml, and 16 µg /ml isoniazid, with and without acetamide, and the plates incubated for 72 h at 37°C.

## RESULTS

### Ca^2+^ imparts structural stability to DesA1

We have earlier observed that DesA1 displays minor changes in fluorescence and far-UV CD spectra upon titration with Ca^2+^ [9]. To investigate metal ion induced structural stability, we performed chemical unfolding of DesA1 with the chaotropic agent urea, in the presence of EGTA and Ca^2+^. Plotting the Trp emission fluorescence of DesA1 as a function of increasing concentrations of urea, the Ca^2+^-induced structural stability with respect to chemical unfolding became apparent (Fig. 1A). We then performed two-state fitting of the data of fraction of native (F_N_) and unfolded (F_U_) DesA1 based on 360/320 ratios (Fig. 1B). The C_1/2_ values of the apo (4 M) and holo (4.5 M) conditions differ by 0.5 M, demonstrating the elevation of Ca^2+^ induced structural stabilisation of DesA1.

**Fig. 1:**
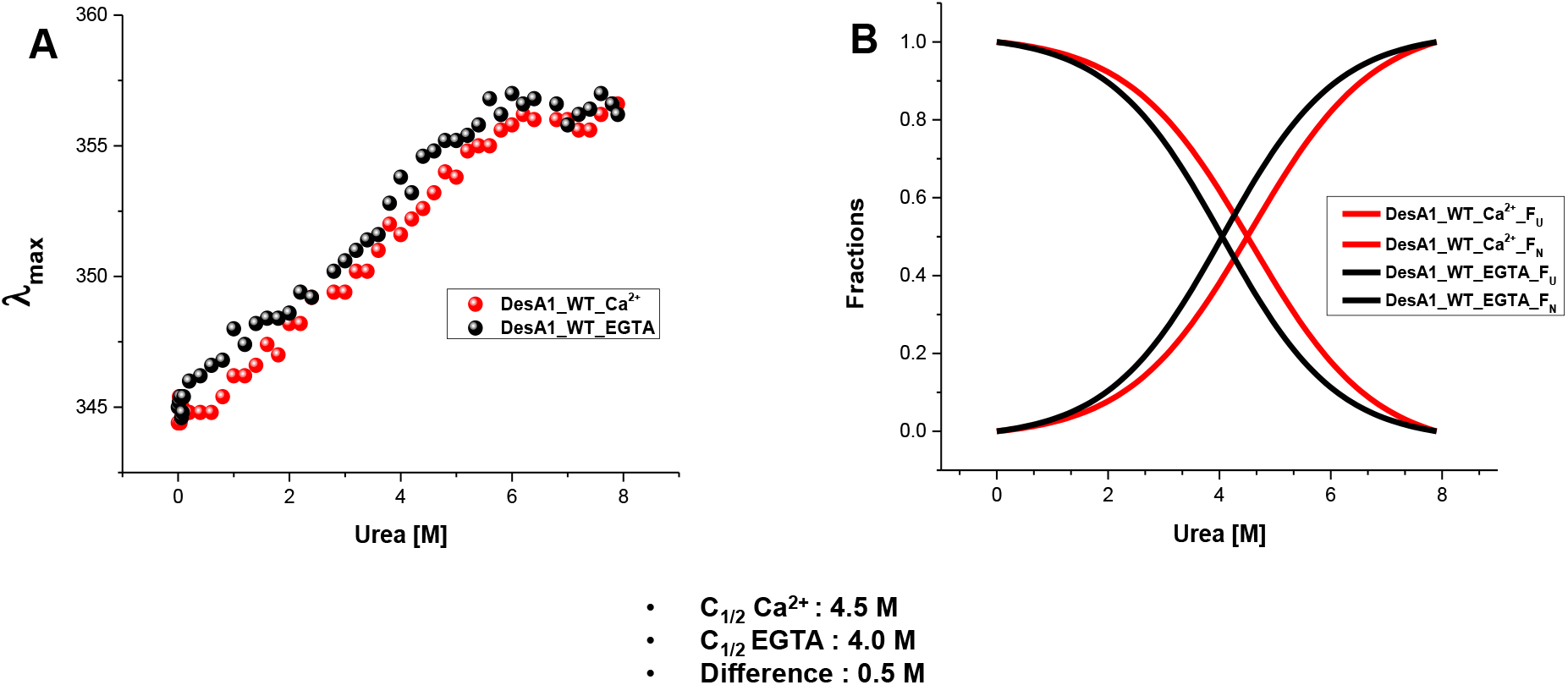
Chemical unfolding of wild type DesA1. A) Trp emission fluorescence spectra of DesA1 as a function of increasing concentrations of urea under Apo (+EGTA) and Holo (+Ca^2+^) conditions. B) Two-state fitting of the fraction of native (F_N_) and unfolded (F_U_) DesA1 based on 360/320 ratios.

### The F303A mutation abolishes the Ca^2+^ binding ability of DesA1

The amino acid sequence of a motif of DesA1 (338aa) which bears the signatures of βγ-crystallin is:269-K**W**RI**F**ERED**F**T**G**EGAKLRDELALVIKDLELACDKFEVSKQRQ-310.The presence of Trp270 prior to conserved Phe273 (WXXF), followed by a Gly at 8^th^ position with a sequence of FXG form a signature of a βγ-crystallins [16]. The position of F303 appears to be conspicuous, and is adjacent to the putative Ca^2+^ co-ordinating residue E304. In an attempt to assess their role, we first introduced Ala in place of Phe (F303A). On Ca^2+^ titration, the intrinsic fluorescence of the F303A mutant did not show any major perturbations (Fig. 2A). Also, far- and near-UV circular dichroism (CD) spectra presented with only trivial changes even at saturable Ca^2+^ concentrations, but without significant Ca^2+^ induced conformational fluctuations (Fig. 2B, 2C). Chemical unfolding analyses of DesA1 F303A in the presence and absence of Ca^2+^ (+EGTA) demonstrate the loss of Ca^2+^-dependent gain of structural stability of DesA1 (Fig. 2D, 2E). Quantitatively, C_1/2_ differs only by 0.01 M: C_1/2_ of the apo form (EGTA was used) was 3.33 M, whereas C_1/2_ of the holo form (in the presence of Ca^2+^) was 3.34 M suggesting a considerable loss of stability due to this mutation. Further, our data of Ca^2+^ binding by isothermal titration calorimetry (ITC) experiments are in agreement with the structural behaviour of the DesA1 F303A mutant, where no measurable heat changes of Ca^2+^ binding were observed (Fig. 2F). These data demonstrate that the F303A mutation significantly compromised the overall gain in structural stability, due to killing the Ca^2+^-binding properties of WT DesA1.

**Fig. 2:**
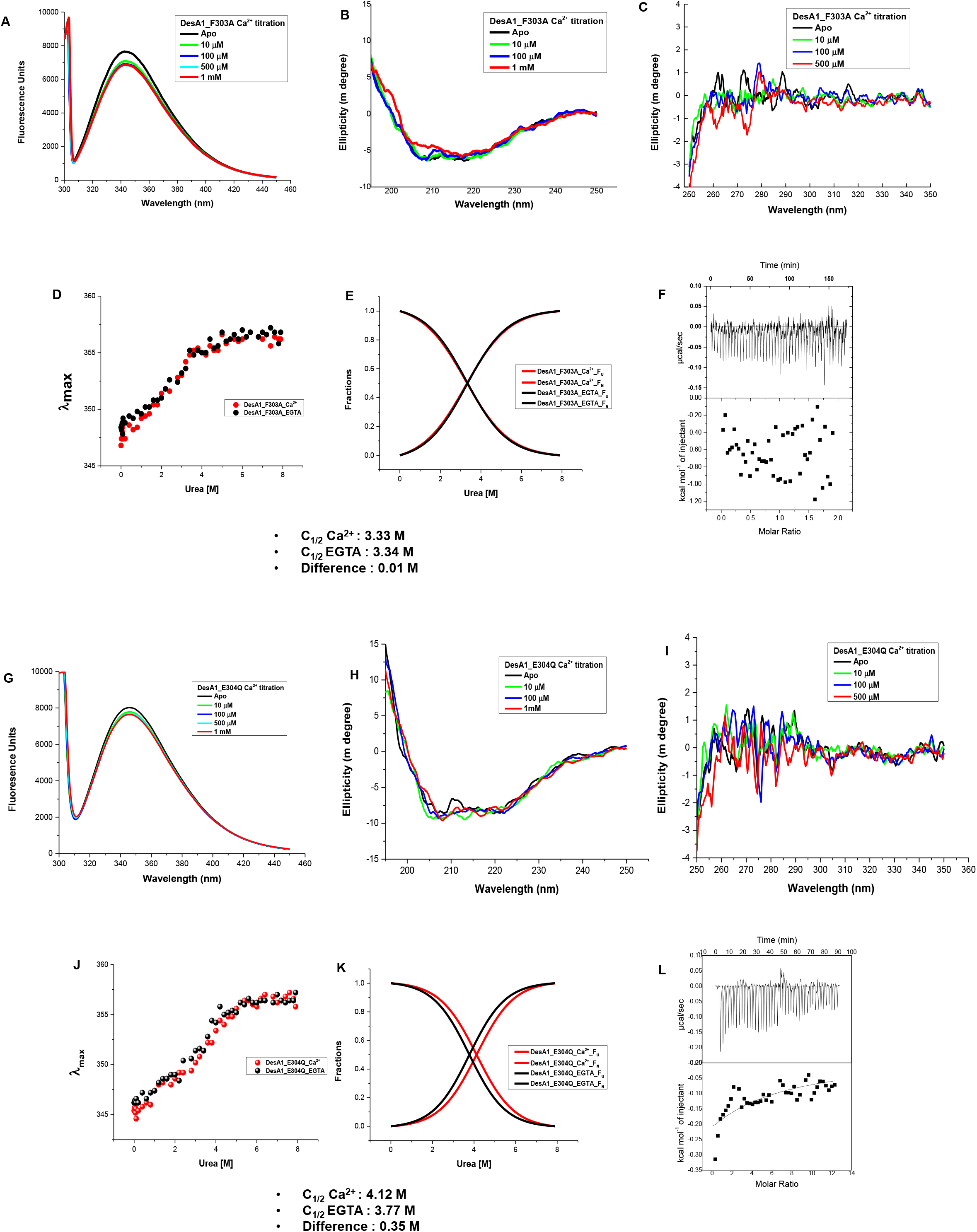
Effect of the F303A and E304Q mutations on structure and Ca^2+^ binding of DesA1. A) Effect of Ca^2+^ titration on intrinsic tryptophan fluorescence spectra of DesA1 F303A. B,C) Far-UV CD spectra of DesA1 F303A to monitor perturbations in the secondary (190/260nm (B)) and tertiary (260/350nm (C)) structures of DesA1 F303A in the presence of 10 μm to 1 mM Calcium Chloride. D)Trp emission fluorescence spectra of DesA1 F303A as a function of increasing concentrations of urea under Apo (+EGTA) and Holo (+Ca^2+^) conditions. E) Two-state fitting of the fraction of native (F_N_) and unfolded (F_U_) protein based on 360/320 ratios. F) Isothermal titration calorigram of Ca^2+^ binding to DesA1 F303A - the concentrations of protein and Ca ^2+^ used in ITC experiments were 50 μM and 5 mM respectively. G) Effect of Ca^2+^ titration on Intrinsic tryptophan fluorescence spectra of DesA1 E304Q H, I) Far-UV CD spectra of DesA1 E304Q to monitor perturbations in the secondary (190/260nm (H)) and tertiary (260/350nm (I)) structures of DesA1E303Q in the presence of 10 μm to 1 mM Calcium Chloride J) Trp emission fluorescence spectra of DesA1 E304Q as a function of increasing concentrations of urea under Apo (+EGTA) and Holo (+Ca^2+^) conditions. K) Two-state fitting of the fraction of native (F_N_) and unfolded (F_U_) protein based on 360/320 ratios. L) Isothermal titration calorigram of Ca^2+^ binding to DesA1 E304A - the concentrations of protein and Ca ^2+^ used in ITC experiments were 50 μM and 5 mM respectively.

### The charge changing mutant E304Q partly retains the properties of DesA1

Having established the criticality of F303 on DesA1 structure and Ca^2+^ binding, we probed the participation of E304 on Ca^2+^ binding and conformation by replacing this Glu residue with a Gln (E304Q). In principle, this mutation should not lead to significant structural changes in the protein, except to alter its negative charge and possibly affect Ca^2+^ binding. The results from fluorescence, far- and near-UV CD spectra indicate an absence of significant structural alterations upon Ca^2+^ titration (Fig. 2A, 2B, 2C). The C_1/2_ values from unfolding experiments (apo 3.77 M and holo 4.12 M) suggest the retention of Ca^2+^-induced structural components (Fig. 2J, 2K). Compared to WT DesA1, the differences in C_1/2_ values of apo and holo proteins, are 0.23 and 0.38 M respectively, whereas the same difference in the case of F303A mutant is 0.7 and 1.2 M. This suggests that the E304Q mutation leads to less structural perturbation in comparison to F303A. The thermogram obtained from ITC experiments, demonstrates a poor but noticeable heat change on Ca^2+^ titration (Fig. 2L), but the data could not be fit into a specific set of sites due to high scatter. These findings suggest that the Ca^2+^-induced structural stability is retained to some extent in DesA1 E304Q compared to F303A.

### Evidence for a compensation mechanism in DesA1 F303A-E304Q

It became evident from the properties of two mutant proteins, that E304 and F303 are crucial for the structure and function of DesA1. To further probe the importance of both residues, we generated a double mutant version of the protein, DesA1 F303A-E304Q. The protein showed minor changes in its fluorescence spectra on Ca^2+^ titration (Fig. 3A), and this titration resulted in significant changes in the far and near-UV CD parameters (Fig. 3B, 3C). These observations were further strengthened by our data from equilibrium unfolding experiments, where the double mutant gained significant stability induced by Ca^2+^, with a difference of C_1/2_ of 0.35 M between the C_1/2,apo_ (3.6 M) and C_1/2,holo_ (3.95 M) forms of the protein, respectively (Fig. 3D, 3E). Compared to DesA1 F303A, the double mutant was structurally somewhat more stable but less than E304Q (Fig. 2D, 2E). Interestingly, ITC measurements showed a significant heat change of binding upon titration with Ca^2+^, and one state fitting model indicated a binding affinity of 289 µM, which is insignificant (global affinity for WT DesA1 is 53 µM, [9]), but with an enhanced heat change compared to E304Q (Fig. 2L). Based on the observations from two single mutants, the double mutant was expected to be structurally extra flexible and functionally compromised; on the contrary, the double mutant appears to show a compensation effect for Ca^2+^ binding when presented together, which may be due to the neutralisation of charge-charge repulsion.

**Fig. 3:**
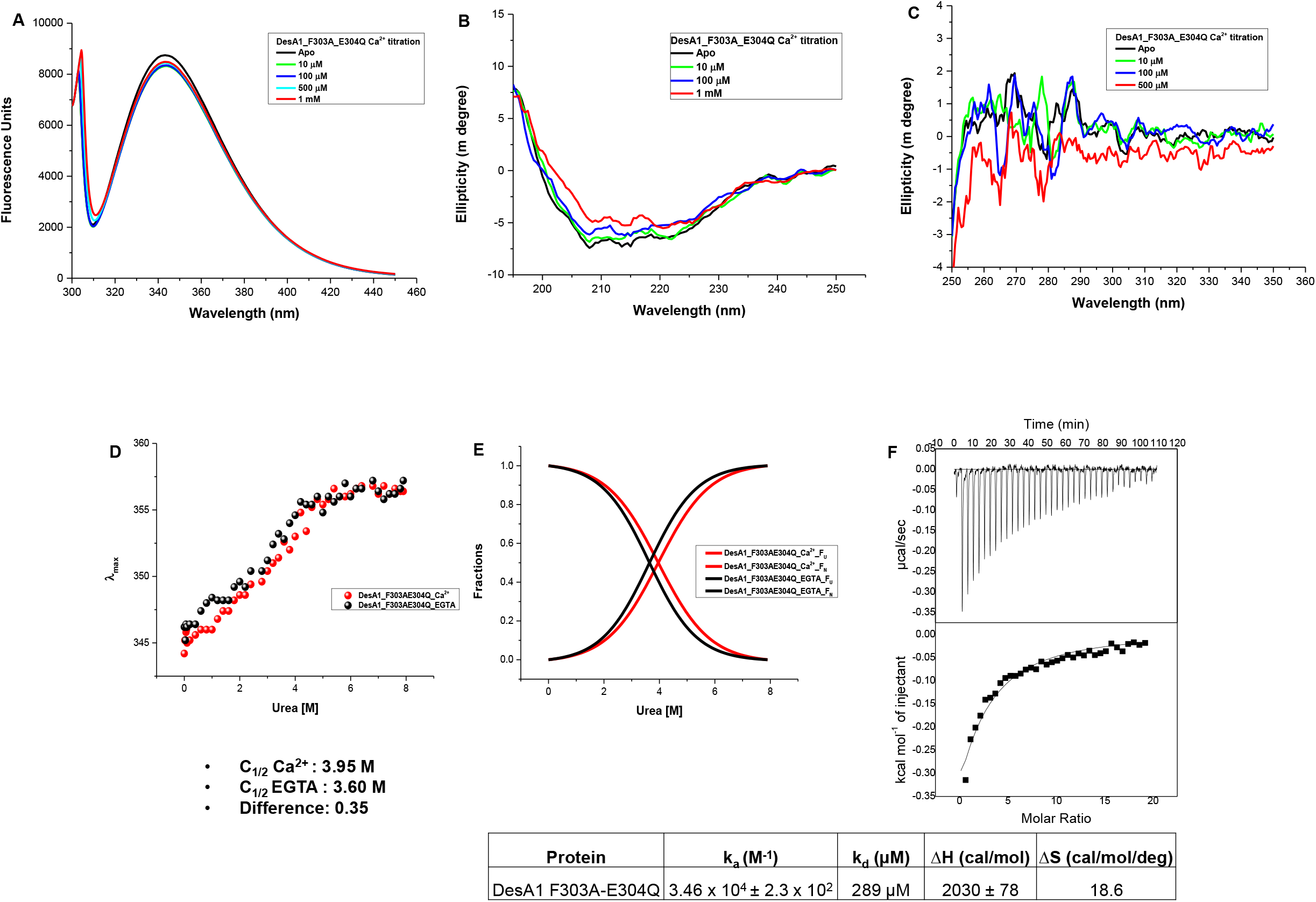
Effect of the F303-E304Q double mutation on structure and Ca ^2+^ binding of DesA1. A) Effect of Ca^2+^ titration on intrinsic tryptophan fluorescence spectra of DesA1 F303-E304Q. B,C) Far-UV CD spectra of DesA1 F303-E304Q to monitor perturbations in the secondary (190/260nm (B) and tertiary (260/350nm (C) structures of DesA1 F303A in the presence of 10 μm to 1 mM Calcium Chloride. D) Trp emission fluorescence spectra of DesA1 F303-E304Q as a function of increasing concentrations of urea under Apo (+EGTA) and Holo (+Ca^2+^) conditions. E) Two-state fitting of the fraction of native (F_N_) and unfolded (F_U_) of the double mutant based on 360/320 ratios. F) Isothermal titration calorigram of Ca^2+^ binding to DesA1 F303-E304Q, the concentrations of protein and Ca^2+^ used in ITC experiments were 50 μM and 5 mM respectively. The table represents the binding kinetics of Ca ^2+^ binding to the double mutant, k_a_-macroscopic association constant, k_d_ - dissociation constant, ΔH - change in enthalpy, ΔS - change in entropy

### F303 is critical for DesA1 function *in vivo*

To assess the implications of the biophysical observations on DesA1 function *in vivo*, a complementation assay was performed using a conditional *M. smegmatis desA1* deletion mutant strain (MSMEG5773) [6], which contains the native gene under the regulation of an acetamide inducible promoter. This strain was transformed with plasmids encoding WT DesA1, DesA1 F303A, DesA1 E304Q, and DesA1 F303A-E304Q, and the ability of each of these to complement the *desA1* defect was tested in the absence of acetamide. Both WT DesA1 and DesA1-E304Q were able to restore the growth phenotype of MSMEG5773, whereas DesA1-F303A, and DesA1-F303A-E304Q were unable to do so (Fig. 4A). This clearly suggests that the F303A mutation results in the complete loss of desaturase activity of DesA1, a reflection of the functional criticality of this residue, and the implicit requirement of Ca^2+^ binding for DesA1 activity.

**Fig. 4:**
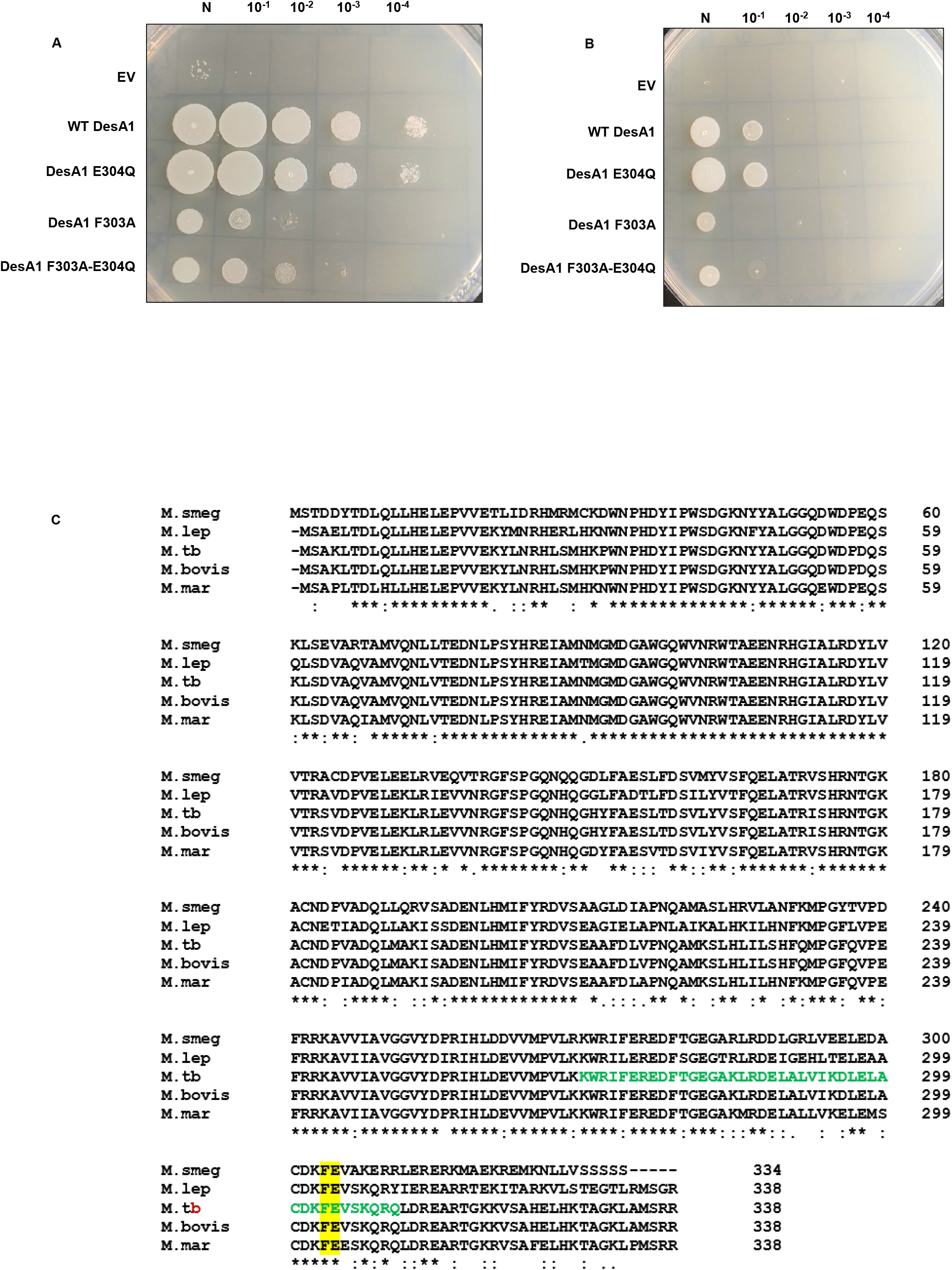
Impact of F303A and E304Q mutations on DesA1 function *in vivo*. A) Complementation assay showing growth profiles of recombinants of a conditional *M. smegmatis desA1* deletion mutant strain, containing plasmids expressing WT *M*.*tb desA1* and its point mutants on Middlebrook 7H10 agar. B) Growth profiles of recombinants of a conditional *M. smegmatis desA1* deletion mutant strain, containing plasmids expressing WT *M*.*tb desA1* and its point mutants, on Middlebrook 7H10 agar containing 4μg/ml Isoniazid. DM - Double mutant, N – Neat. C) Multiple Sequence Alignment of DesA1 homologs from *Mycobacterium smegmatis* (M. smeg), *Mycobacterium leprae* (M. lep), *Mycobacterium tuberculosis* (M. tb), *Mycobacterium bovis* (M. bovis), and *Mycobacterium marinum* (M. mar) performed using Clustal Omega (https://www.ebi.ac.uk/Tools/msa/clustalo/). The sequence in green corresponds to the βγ-crystallin motif in *M. tb* DesA1, the conserved residues corresponding to F303 and E304 in *M. tb* DesA1 are highlighted in yellow.

### Dysfunction of DesA1 leads to compromised permeability in *M. smegmatis*

Since a loss in DesA1 function leads to compromised mycolic acid synthesis and altered integrity of the mycobacterial cell envelope, we examined the permeability of recombinant MSMEG5773 transformed with plasmids encoding WT DesA1, DesA1 F303A, DesA1 E304Q, and DesA1 F303A-E304Q, using sensitivity to the first line anti-TB drug Isoniazid as a surrogate marker. Under non-permissive conditions, the recombinants expressing DesA1 F303A, and DesA1 F303A-E304Q displayed enhanced susceptibility to 4 µg/ml isoniazid (Fig. 4B) indicative of their increased permeability due to the loss of DesA1 function, in comparison to those expressing WT DesA1 and the E304Q mutant.

## Discussion

βγ-Crystallins are well-established Ca^2+^-binding proteins with a wide distribution across different life forms [17-19]. We have previously identified a βγ-crystallin type motif in the sequence of *M. tb* DesA1 and demonstrated that, like many other microbial homologues, that it also binds Ca^2+^ (9). Yet, the obvious question that crossed our minds was the functional significance of this motif in DesA1. Based on our knowledge of numerous βγ-crystallins, other than Ca^2+^ binding, we did not hesitate in proposing that a βγ-crystallin motif would provide structural stability to DesA1. In this study, we present some novel features of DesA1, including the property that Ca^2+^ binding imparts structural stability to this protein. The role of a phenylalanine residue (F303) in the βγ-crystallin signature sequence was also identified as a hot point implicated in Ca^2+^ binding. Mutating it (to an Ala) causes a significant loss of overall protein stability, and a complete loss of Ca^2+^ binding attributes of DesA1 (Fig 2D, 2E, 2F). In the absence of the availability of its crystal structure, no information regarding the coordination chemistry is available; hence, the serendipitous finding of the F303 residue controlling Ca^2+^ binding proved to be significant.

This mutant (F303A) provided us with the ideal opportunity to exploit it as a probe to investigate the role of Ca^2+^ with respect to DesA1 in *M. tb*. By comparing DesA1 (which binds Ca^2+^) with DesA1-F303A (which does not bind Ca^2+^), we were able to strategically examine the functional importance of Ca^2+^ binding. As seen by our data, the Ca^2+^ disabled mutant (DesA1-F303A) failed to complement the growth phenotype or led to a significant compromise in complementation of a conditionally complemented *desA1* deletion mutant strain of *Mycobacterium smegmatis*. Consequently, in brief, the defective and impaired desaturase activity and enhanced permeability, observed in the F303A and F303A-E306Q mutants, emphasised the significance of the involvement of Ca^2+^ *via* binding to the βγ-crystallin motif of DesA1 (Fig. 4A, 4B). Multiple sequence alignment shows that this region of the protein sequence is well conserved in all mycobacterial DesA1 homologs, including absolute conservation of the residues corresponding to F303 and E304 of *M. tb* DesA1 (Fig. 4C). It is therefore highly likely that the essential Ca^2+^ dependent function of this protein is conserved across mycobacteria.

The role of Ca^2+^ in controlling virulence is an important aspect required for therapeutic control [20, 21]. The presence of a Ca^2+^-binding βγ-crystallin motif in selective microbes, including those that are pathogenic, points to its involvement in virulence [22]. Our study proves that Ca^2+^ binding is essential for DesA1 function and suggests that the structural changes induced by Ca^2+^ binding lead to increased stability of the protein. Being a modest affinity Ca^2+^ binding protein, DesA1 is not a Ca^2+^ sensor. The role of Ca^2+^ therefore in DesA1 function, became much more obvious in the complementation and membrane permeability experiments, where the complete loss of Ca^2+^ binding (in DesA1 F303A) and low affinity Ca^2+^ binding (in DesA1 F303A-E306Q), led to dysfunction of DesA1. Additionally, this work is a presentation of the structure-function relationship of the βγ-crystallin motif of DesA1, and highlights the significance of βγ-crystallin recruitment in *M. tb*. The increased susceptibility of DesA1 mutants to isoniazid (and perhaps to other anti-TB drugs as well) (Fig 4B) taken together with the essentiality of DesA1 in *M. tb* (6), makes this protein an attractive target for the development of anti-tubercular drugs based on Ca^2+^ chelation.

## Acknowledgement

We are grateful to Dr. Apoorva Bhatt, University of Birmingham, UK for the kind gift of pMV206-Apra, an *E. coli*-Mycobacterium shuttle vector, and *M. smegmatis* ΔMSMEG5773, the conditional knockout strain of *M. smegmatis desA1*

## Author contributions

T.R.R., M.S.S and R.P.M designed the study. M.S.S, R.P.M, U.K., C.V.Y. and N.N. performed the experiments. T.R.R., M.S.S, R.P.M and U.K. analysed the data. T.R.R., M.S.S, R.P.M, U.K. and Y.S. wrote the manuscript.

