## Supplementary material for "Dissecting the Ca^2+^ dependence of *Mycobacterium tuberculosis* DesA1 function": Table S1

**Table S1: Primers used in this study**

| | Name | Primer Sequence |
|---|---|---|
| 1 | F3030A-DesA1-FP | 5'-TGCGACAAGGCCGAGGTGTCCAAG-3' |
| 2 | F303A-DesA1-RP | 5'-CTTGGACACCTCGGCCTTGTCGCA-3' |
| 3 | E304Q-DesA1-FP | 5'-TGCGACAAGTTCCAGGTGTCCAAG-3' |
| 4 | E304Q-DesA1-RP | 5'-CTTGGACACCTGGAACTTGTCGCA-3' |
| 5 | F3030AE304Q-DesA1-FP | 5'-TGCGACAAGGCCCAGGTGTCCAAG-3' |
| 6 | F3030AE304Q-DesA1-RP | 5'-CTTGGACACCTGGGCCTTGTCGCA-3' |
| 7 | pMV261-DesA1-FP | 5'-ATCGAGGATCCATGTCAGCCAAGCTGACCG-3' |
| 8 | pMV261-DesA1-RP | 5'-TGCATGAATTCCTAACGACGGCTCATCGC-3' |
